# The short-sequence design of human chromosomes

**DOI:** 10.1101/365262

**Authors:** Guillermo Lamolle, Victor Sabia, Héctor Musto, Giorgio Bernardi

## Abstract

Recent investigations have shown that isochores are characterized by a 3-D structure which is primarily responsible for the topology of chromatin domains. More precisely, an analysis of human chromosome 21 demonstrated that GC-poor isochores are low-heterogeneity sequences characterized by the presence of oligo-Adenines that are intrinsically stiff, curved and unfavorable for nucleosome binding. This leads to a structure of the corresponding chromatin domains, the Lamina Associated Domains, or LADs, which is well suited for interaction with lamina. In contrast, the high-heteorogeneity GC-rich isochores are in the form of compositional peaks characterized by gradients of oligo-Guanines that lead to increasing nucleosome depletions in the corresponding chromatin domains, the Topological Associating Domains, or TADs. These results encouraged us to investigate in detail the di- and tri-nucleotide profiles of 100Kb segments of chromosome 21, as well as those of the di- to octa-Adenines and di- to octa-Guanines in several regions of the chromosome. The results obtained show that the 3-D structures of isochores and chromatin domains depend not only upon oligo-Adenines and oligo-Guanines but also, to a lower but definite extent, upon the majority of di- and tri-nucleotides. This conclusion, which applies to all human chromosome, has strong implications for the biological role of non-coding sequences.

---

The human genome (like other mammalian genomes) is compositionally compartmentalized into isochores, large (≥ 0.2 Mb), “fairly homogeneous” DNA sequences that belong to five families, L1, L2, H1, H2, H3, characterized by increasing GC ranges, increasing compositional heterogeneities and increasing gene densities^1–3^ (see Supplementary Table S1, Fig. S1 of ref.4 and ref.5 for a review). While the functional importance of isochores was evident for a long time because of their correlations with all genome properties that were tested and led to their definition as “a fundamental level of genome organization”^6^, the basic reason for their existence is not yet understood^6^, even if links with chromatin structure^1^ and with nucleosome positioning and density^7^ were predicted.

Recent investigations^8^ showed that maps of GC-rich and GC-poor isochores of all human and mouse chromosomes match maps of TADs (0.2-2 Mb in size^9^) and maps of LADs (~ 0.5 Mb medium size^10^), respectively. Moreover, the average size of human isochores, ~0.9 Mb, is very close to that, ~0.88 Mb, of 91% of the mouse chromatin domains^11^. Finally, both chromatin domains and isochores are evolutionarily conserved in mammals^2,11^.

Very recent work^4^ led to the discovery that, topologically, isochores are characterized by 3-D structures that play a crucial role in the formation of TADs and LADs and that the topology of isochores depends upon some of their short sequences, such as oligo-As and oligo-Gs. Here, the wider problem concerning the general involvement of oligonucleotides in the 3D structure of the genome was approached.

## RESULTS

### The GC profile of chromosome 21 and its connection with chromatin domains

The profile of chromosome 21, as obtained using a point-by-point plot of GC levels of 100 Kb blocks (Fig.1A,B), is shown as a reference profile. Indeed, this profile (split into several regions, numbered 1 to 6) provided the first suggestion of a 3-D structure of isochores by showing a low-heterogeneity L1 region 2 and a series of single H1 (a) and L2^+^ peaks (b to f) in regions 1 and 3, two multi-peak regions 4 and 5 corresponding to H1 and H2 isochores, respectively, and a region 6 characterized by very sharp peaks ranging from 40-45 *%* to 55-60% GC. It has been recently shown^4^ that, while the GC-poor, low-heterogeneity L1 region (as well as L2^-^ regions only represented in chromosome 21 by the “valley” isochore Y) corresponds to LADs, the GC-rich, high-heterogeneity L2^+^, H1, H2, H3 peaks corresponds to TADs.

**Figure 1.**
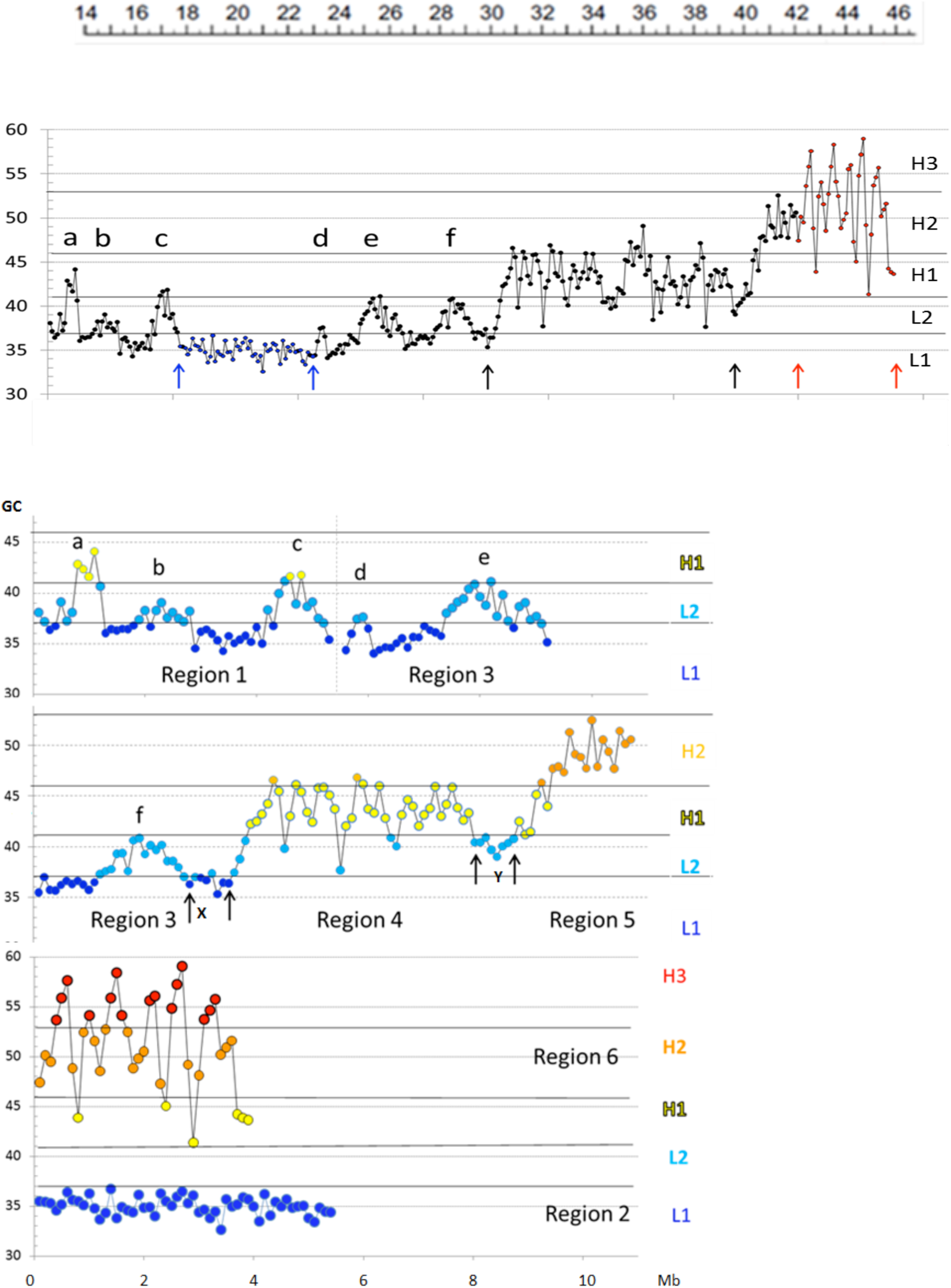
A. Compositional profile of human chromosome 21. (from ref.4). GC levels of 100Kb windows are presented in a point-by-point plot. This figure shows that: region 1 comprises a (rare) H1 single peak (a) and two L2^+^ peaks (b, c); region 2 corresponds to a compositionally flat L1 isochore; region 3 comprises three L2^+^ peaks (d, e, f); regions 4 and 5 comprise the multi-peak H1 and H2 isochores, respectively; region 6 comprises H3 peaks. Black, blue and red lines, as well as arrows (X,Y), separate regions 1 to 6. **B. Panel A is displayed at a higher magnification** (from ref.4). DNA sequences from isochore families L1 to H3 are represented in different colors, deep blue, light blue, yellow, orange, red, respectively. Double lines separate two “valley” L2^-^ isochores, X and Y. Permission to publish Fig.1 was obtained from the copyright owner.

### The di-nucleotides of chromosome 21

An analysis^4^ of previous results^7,12^ (see Supplementary Fig.1) showed that “A/T-only” and “G/C-only” di-nucleotides were the most abundant ones in GC-poor and GC-rich isochores, respectively (with AA, TT and GG, CC, respectively, reaching the highest levels); in contrast, the “mixed” dinucleotides (comprising both A and T or G and C) showed only modest differences in different isochore families. For these reasons, the profiles of the three sets of di-nucleotides were investigated separately.

The “G/C-only” di-nucleotides showed profiles (Fig.2A) that were similar to GC profiles (Fig.1). These profiles were characterized by a low-heterogeneity region 2 (with a ~4% level), a series of peaks c, d, e, f, in regions 1 and 3, two multi-peak regions 4 and 5 and a series of sharp GpG and CpC peaks in region 6. GpC showed a slightly lower level in region 2 and peaks covering a smaller range in region 6 compared to GpG and CpC. CpG showed the lowest levels (~1% in region 2 and small peaks in region 6), as expected from the well-known shortage of this di-nucleotide^13^.

**Figure 2.**
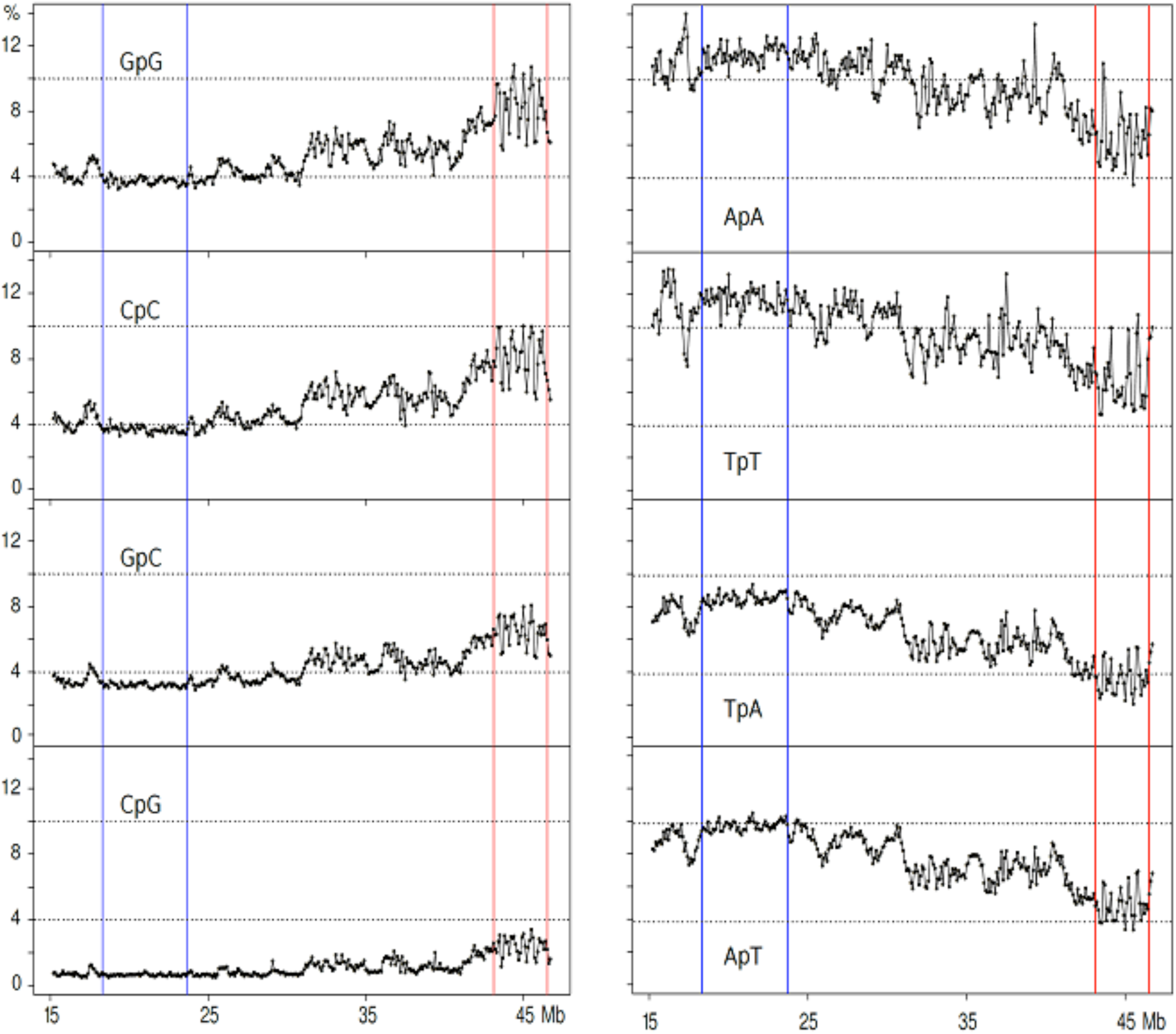
Profiles of (A) “G/C-only”, and (B) “A/T-only” di-nucleotides. Please note that this and the following profiles lack the centromeric region comprising the a and b peaks of Fig.1, because derived from a different sequence release (see Methods). Blue and red lines define regions 2 and 6, respectively. Black lines separate region 4 from the two contiguous regions 3 and 5.

In contrast, the “A/T-only” di-nucleotides (Fig.2B) showed much higher levels in region 2 compared to the “G/C only” di-nucleotides (~12% for ApA and TpT, with lower values, 10% and 9%, for ApT and TpA, respectively). In region 6, ApA and TpT showed sharp peaks although reaching lower levels compared to “G/C-only” di-nucleotides (these levels being even lower in the case of ApT and TpA) and higher levels and sharper peaks in regions 1 and 3-5, again especially in the case of ApA and TpT. In the case of region 1, the A/T peaks preceded the G/C peak c. Likewise, the A/T peaks of region 3 alternated with the G/C peaks e and f. In region 5, the downward A/T slope corresponded to an upward G/C slope. The more complex region 4 will be presented later.

“Mixed di-nucleotides”, namely di-nucleotides comprising both A/T’s and G/C’s, showed flat profiles with 5-7% values in region 2. Minute peaks were present in region 6 in the case of GpT, ApC, CpA and TpG (Fig. 3A and 3B).

**Figure 3.**
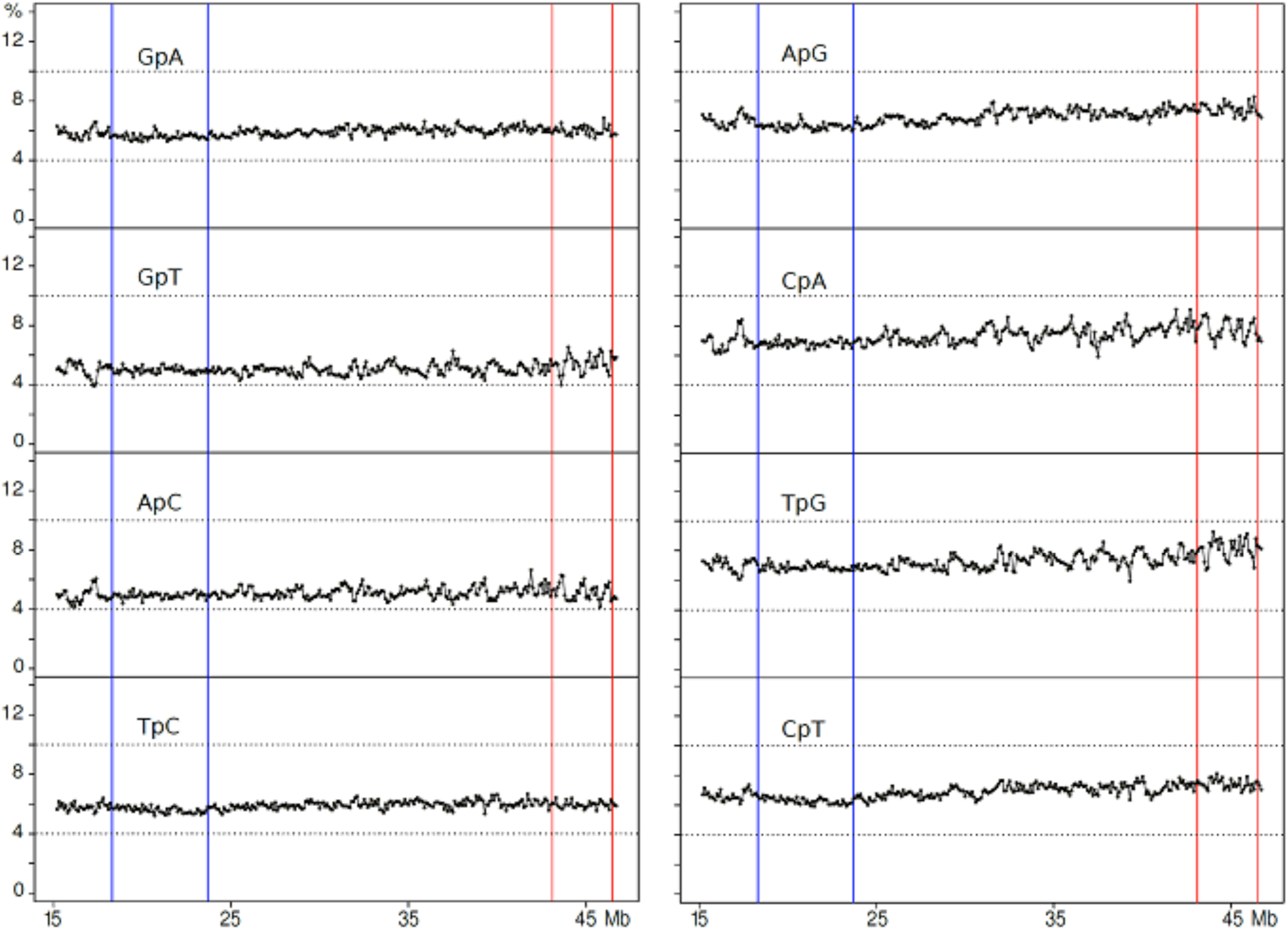
Profiles of “mixed” di-nucleotides.

### "G/C-only” and “A/T-only” tri-nucleotides

Supplementary Fig.S2 presents previous results^12^ to show that tri-nucleotides, like di-nucleotides, are characterized by frequencies in different isochore family that are different for “GC-only”, “AT-only” and “mixed” tri-nucleotides.

Among the “G/C-only” tri-nucleotides, the GGG and CCC profiles were sharper than the GGC and GCC profiles and were similar to GC profiles in which one could detect the single c to f peaks (including the minute d peak just on the right side of the second blue line), the sharp peaks of region 6 and the flat low-level (~1%) region 2 (Fig.4A). In contrast, the “G/C-only” tri-nucleotides that comprised the CG doublet showed, as expected, very flat, very low profiles, as well as minute peaks in region 6 (Fig. 4B).

**Figure 4.**
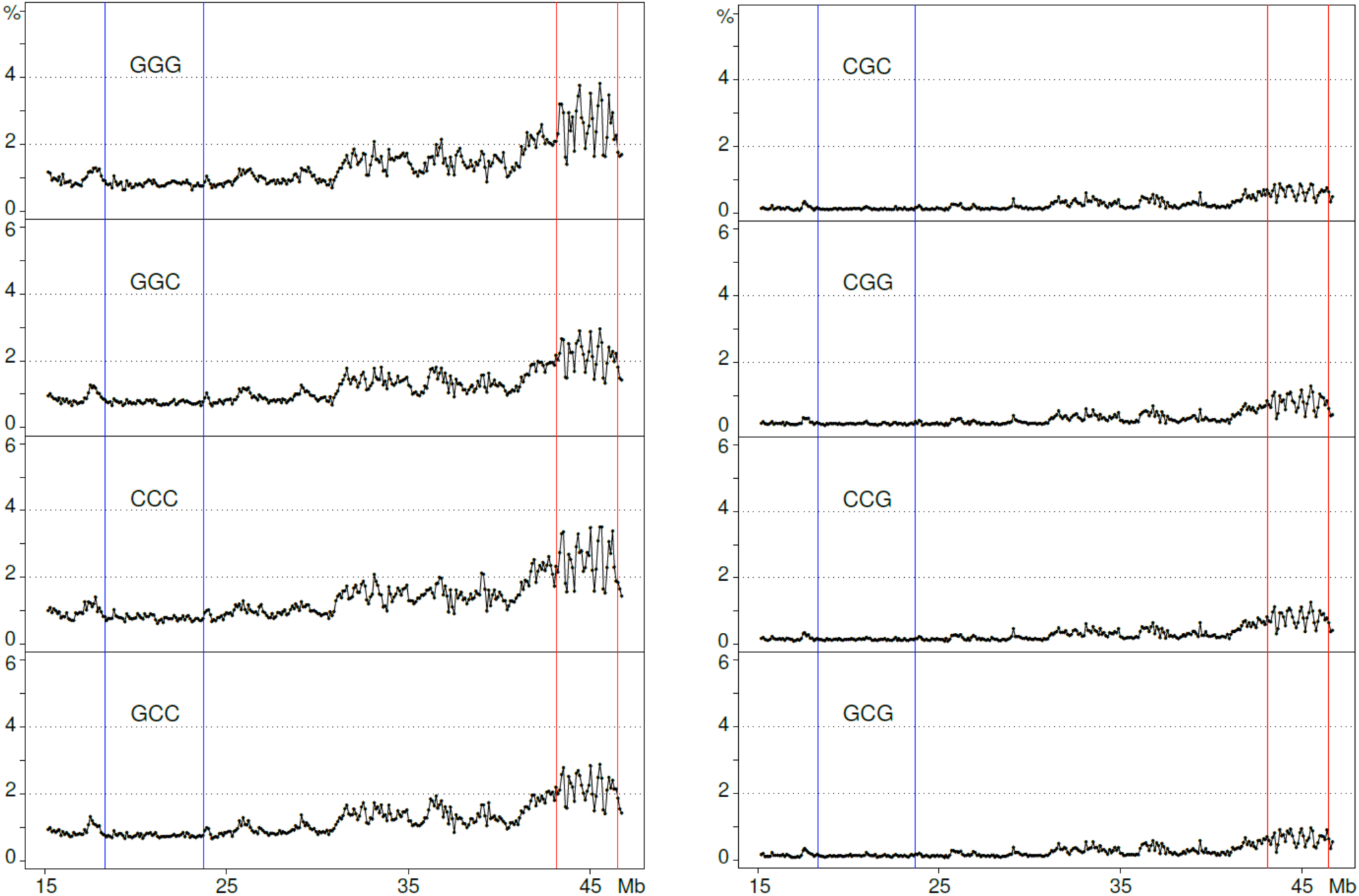
**Profiles of “G/C-only” tri-nucleotides** not comprising (**A**) or comprising (**B**) the CpG doublet.

Among “A/T-only” tri-nucleotides, AAA and TTT showed very high levels (~5%) in region 2 and a series of peaks in regions 1 and 3-6, the peaks being extremely sharp in region 6 (Fig.5A), whereas the other profiles (*e.g.*, ATA) were lower in region 2 (~3%) and covered a smaller range in region 6 (Fig.5B). Moreover, as in the case of dinucleotides, all A/T peaks of region 1 preceded peak c (that corresponds to a trough in the profile). The apparent contradiction of finding both “G/C-only” and “A/T-only” in region 6 (as well as in regions 1 and 3 to 5) could be solved by comparing in detail the two series of profiles from di- to octa-nucleotides for regions 2, 6 and 4.

**Figure 5.**
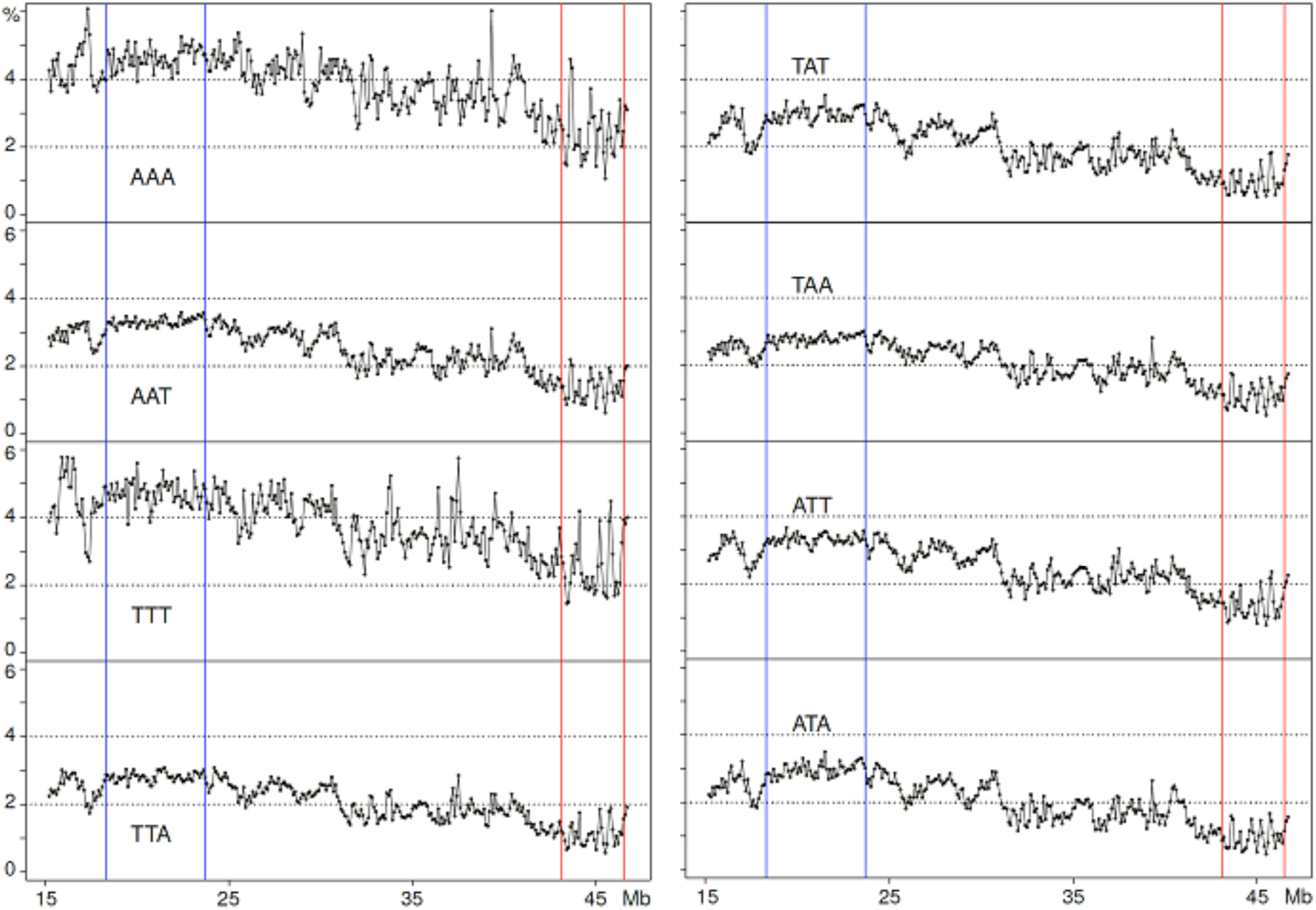
Profiles of “A/T-only” tri-nucleotides.

### A comparison of di- to octa-As and Gs from regions 2, 6 and 4

The results obtained for di-, tetra- and octa-A’s and G’s for regions 2 and 6 are presented in Figs. 6A, 6B, those for intermediate oligonucleotide sizes are shown in Supplementary Figs.S3. In region 2, the profiles are characterized, as expected, by high levels of oligo-A’s, their increasing sizes (from 2 to 8 nucleotides) having an increasingly higher level compared to the same-size oligo-G’s. In region 6, the di-nucleotides plots showed higher levels for GpG compared to ApA, a ratio which, however, progressively decreased with increasing oligonucleotide size. The most important finding was, however, that the oligo-A and oligo-G peaks were alternating, the former ones corresponding to the troughs of the latter ones. Very interestingly, the alternation was also found in the complex, multipeak region 4 (Fig.7).

**Figure 6.**
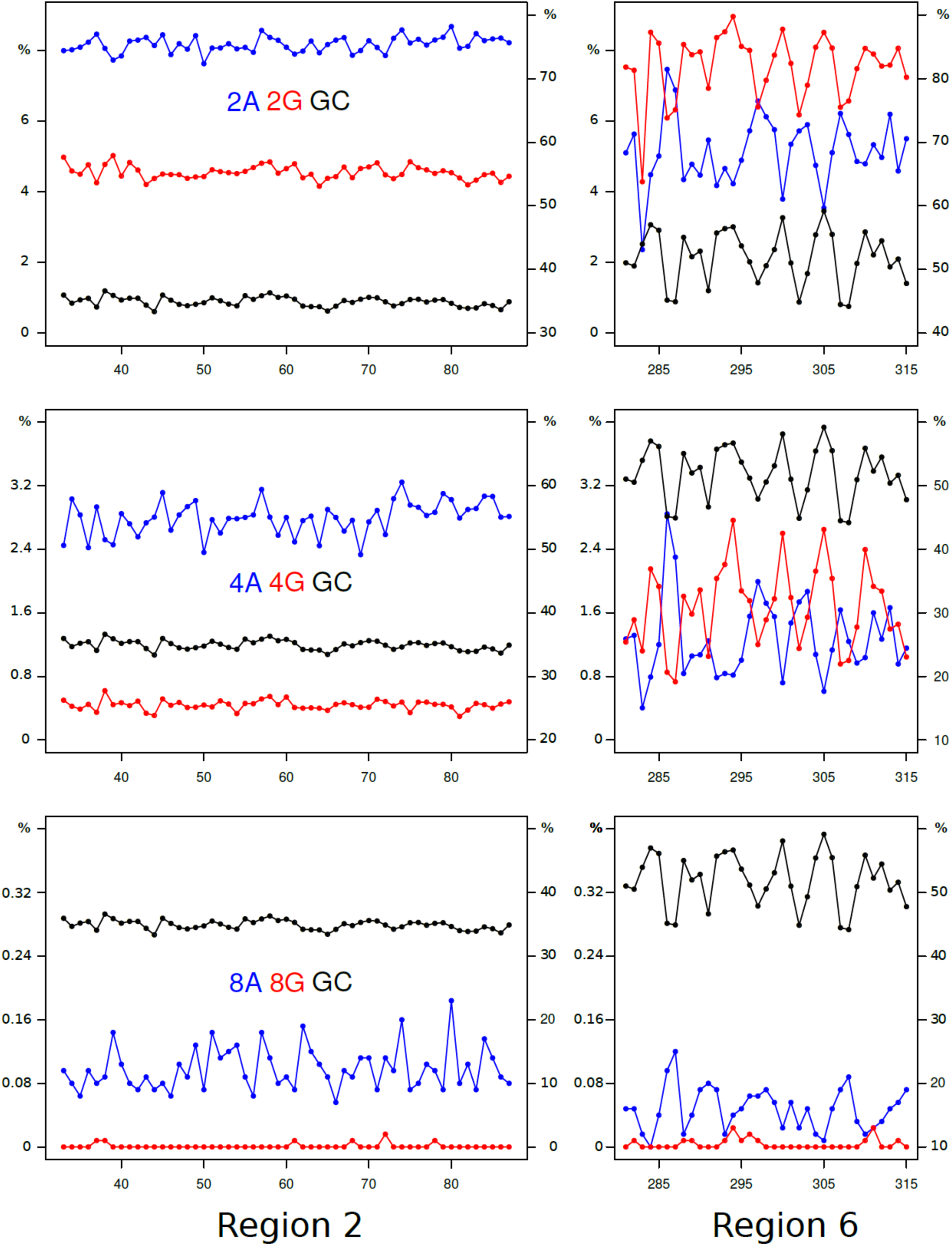
Profiles of di- tetra- octa- As. (blue plots) **and Gs** (red plots; left scales) and GC profiles (black plots; right scales) are displayed for regions 2 and 6.

**Figure 7.**
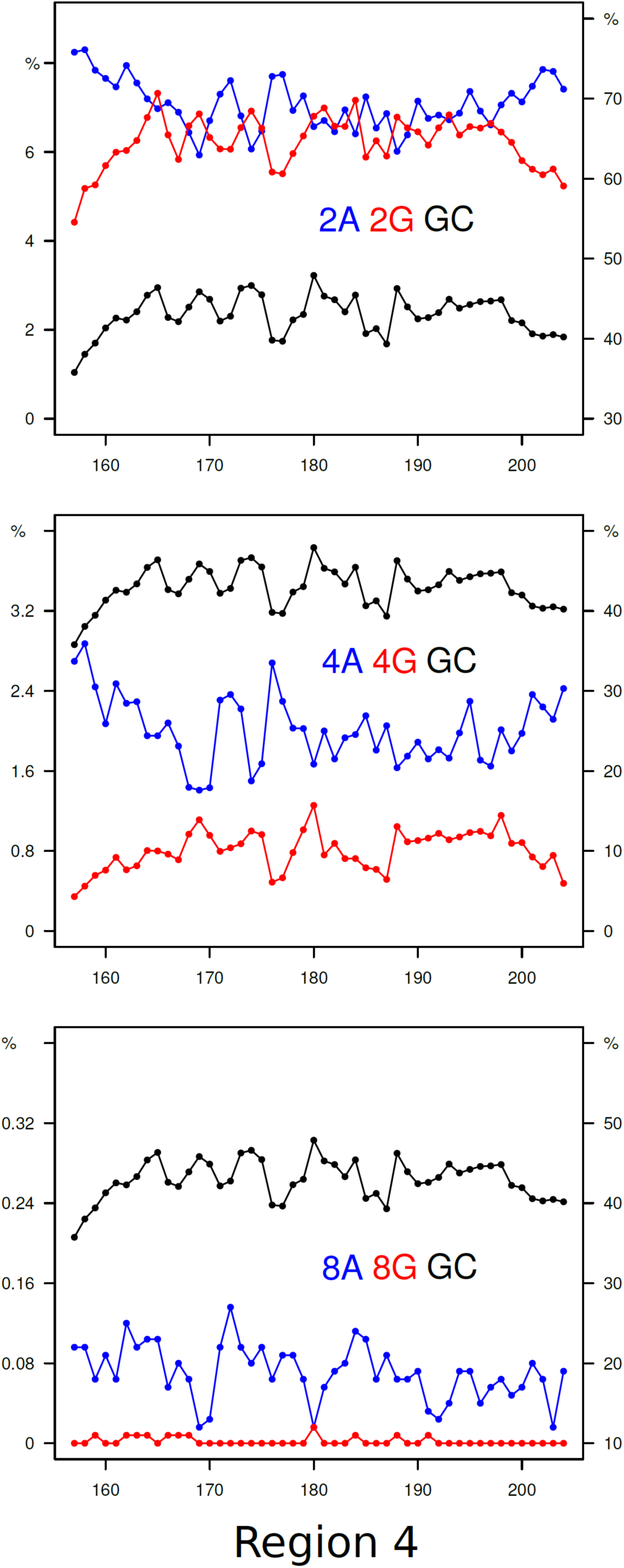
**Profiles of di- tetra- and octa- As and Gs** are displayed for region 4. Scale indications as in Fig.6.

### “Mixed” tri-nucleotides

The “mixed” tri-nucleotides were shown to behave in a slightly different way according to whether they comprised two G or C and one A or T or *viceversa*, two A or T and one G or C. Indeed, the first set (see Supplementary Figs.S4A and S4B) showed low values (~1-2%) in region 2 and small peaks (or, rarely, no peaks) in region 6. (As expected, tri-nucleotides comprising the CpG doublet showed extremely low values in region 2 and no peaks in region 6). The second set showed practically even values (~2% or only slightly lower values) over the whole profile, only in some cases showing minute peaks in region 6 (see Supplementary Figs. S5A and S5B).

## DISCUSSION AND CONCLUSIONS

The first question to be discussed is whether the results presented here for the di- and tri-nucleotides of chromosome 21 are generally valid for the whole human genome. This point is definitely supported by a comparison of the present results with those previously obtained for the corresponding isochore families of the human genomes^7,12^.

In the case of “G/C-only” di-nucleotides, they are at a ~4% level in L1 region 2 (with the exception of the CpG values that are much lower) when peaks in H3 region 6 reach 10% for GpG and CpC. These results are in agreement with the corresponding values of L1 and H3 isochores from the whole human genome. In the case of the “AT-only” di-nucleotides, they amount to 10-12% in L1 region 2, again to the same values found for the L1 isochore family from the whole genome. If one looks at this point in more detail, one can see that the slightly lower value of ApT (~10%) and the still lower value of TpA (~9%) found for region 2 also match the corresponding values for the whole genome (see Supplementary Fig.S1). All these dinucleotides show a series of peaks in region 6 that cover a 4% to 10% range for ApA and TpT, but decrease to 4-6% for ApT and 3-5% for TpA; similar trends are shown in regions 1 and 3 to 5.

In the case of “G/C-only” tri-nucleotides, they show low, flat profiles (~1%) in L1 region 2 and peaks in H3 region 6 (with much lower values if comprising CpG doublets), as well as in region 1 and 3-5; these results are comparable with those of the whole genome (see Supplementary Fig.S2). “A/T-only” tri-nucleotides showed high values, 3-5%, in region 2 and smaller peaks in region 6, once more in agreement with the whole genome results. The slightly different behaviors of “mixed” tri-nucleotides according to whether they have 2 A/T and 1 G/C or *viceversa*, was also found in previous work on the whole genome^12^ (see Supplementary Fig.S2).

The main conclusion, however, of the present work concerns the involvement of di- and tri-nucleotides in the topology of isochores. Indeed, “GC-only” di- and tri-nucleotides showed peaks in region 6 and low, flat levels in region 2 (the special case of CpG doublets in di- and tri-nucleotides was already mentioned). In the case of “A/T-only” di- and tri-nucleotides, levels were high in region 2, whereas peaks covering a lower range compared to “G/C-only” di- and tri-nucleotides were present in region 6 (see below). “Mixed” tri-nucleotides showed more distinct peaks in region 6 in the case of 2 G/C compared to that of 2 A/T.

The detailed analysis of oligo-A’s and oligo-G’s in region 2 revealed that the A/G ratio shows a very strong increase from di-nucleotides to tetra-nucleotides and octa-nucleotides. In the case of region 6, the profiles are characterized by oligo-A and oligo-G peaks that alternate and show a G/A ratio decreasing from di- to tetra- and octa-nucleotides, a remarkable result in view of the high GC level of region 6. This is a very important point which also explains the results of “A/T-only” and “G/C-only” peaks in regions 1 and 3 to 5, the case of region 4 having been investigated in detail. Summing up, interspersed “G/C-only” di- and tri-nucleotides and oligo-Gs are distributed in increasing gradients in the peaks, whereas they have a low flat distribution in L1 region 2. In contrast, interspersed “A/T-only” di- and tri-nucleotides and oligo-As are distributed in increasing gradients in peaks that alternate with the G/C peaks, and have high levels in the L1 region 2.

One should now consider, as already pointed out^4^, that oligo-A’s and oligo-G’s are intrinsically stiff for different structural reasons^14^; more specifically, oligo-A’s are “intrinsically curved”^15^. Two different classes of isochores can, therefore, be distinguished from a topological viewpoint: 1) the L1 isochores that are compositionally even yet locally stiff because of the presence of interspersed oligo A’s and 2) the L2, H1, H2 and H3 isochores that are characterized by increasing densities of interspersed oligo-G’s alternating with increasing densities of interspersed oligo-A’s.

A crucial point is now to consider that the features just described for the topology of GC-poor and GC-rich isochores, namely the presence of interspersed oligo-As and oligo-Gs, are strongly inhibitory to nucleosome formation^14^ and account for the different structures of chromatin domains, LADs and TADs, as proposed elsewhere^4^. Two final consideration are the following. In the case of LADs (*e.g.*, the L1 sequence of region 2), all di- and tri-nucleotide profiles are compositionally flat even if characterized by different levels. In contrast, in the case of TADs, three different situations exist according to whether di- and tri-nucleotide profiles: 1) follow the GC profiles of GC-rich isochores and, obviously contribute to them; 2) are in the form of “minute peaks”, contributing less to the GC profiles; and 3) are flat, not contributing to the GC profiles. In other words, not only oligo-A’s and oligo-G’s are involved in isochore and chromatin topology, but, also, to a lower yet definite extent, all “GC-only” and “AT-only”, as well as a large number of “mixed” di- and tri-nucleotides, as indicated by their gradients in peak regions. This opens the door to a relevant role of short sequences that represent a very sizable fraction of the ~98% non-coding DNA present in human genome.

## Acknowledgements

Guillermo Lamolle thanks Dr. Takashi Gojobori and Dr. Toyonori Sakata for a Gojobori-Sakata fellowship which allowed him to visit Giorgio Bernardi for discussing the present work.

Giorgio Bernardi thanks Paolo Ascenzi for hospitality and Marta Ritucci for excellent technical help.

## AUTHORS’ CONTRIBUTIONS

G. L. and V.S. did the work under the guidance of H. M. and G.B. G. B. wrote the paper.

## METHODS

The data of the sequences that were used for the analyses are the following: gi|568815577|ref|NC_000021.9| Homo sapiens chromosome 21, GRCh38.p7 Primary Assembly Genome Reference Consortium Human Build 38 patch **release 7** (GRCh38.p7)

